# pH selects for distinct N_2_O-reducing microbiomes in tropical soil microcosms

**DOI:** 10.1101/2023.11.29.569236

**Authors:** Yanchen Sun, Yongchao Yin, Guang He, Gyuhyon Cha, Héctor L. Ayala-del-Río, Grizelle González, Konstantinos T. Konstantinidis, Frank E. Löffler

## Abstract

Nitrous oxide (N_2_O), a greenhouse gas with ozone destruction potential, is mitigated by the microbial reduction to dinitrogen catalyzed by N_2_O reductase (NosZ). Bacteria with NosZ activity have been studied at circumneutral pH but the microbiology of low pH N_2_O reduction has remained elusive. Acidic (pH<5) tropical forest soils were collected in the Luquillo Experimental Forest in Puerto Rico, and microcosms maintained with low (0.02mM) and high (2mM) N_2_O assessed N_2_O reduction at pH 4.5 and 7.3. All microcosms consumed N_2_O, but long lag times of up to 7 months were observed in microcosms with 2 mM N_2_O. Comparative metagenome analysis revealed that *Rhodocyclaceae* dominated in circumneutral microcosms under both N_2_O feeding regimes. In acidic microcosms, *Peptococcaceae* dominated in high-N_2_O, and *Hyphomicrobiaceae* in low-N_2_O microcosms. Seventeen metagenome-assembled genomes (MAGs) recovered from these microcosms harbored *nos* operons, with all eight MAGs derived from acidic microcosms carrying the clade II type *nosZ*, lacking nitrite reductase genes (*nirS*, *nirK*). Five of these MAGs represented novel taxa indicating an unexplored N_2_O-reducing diversity exists in acidic tropical soils. A survey of pH 3.5-5.7 soil metagenome datasets revealed that *nosZ* genes commonly occur, suggesting broad distribution of N_2_O reduction potential in acidic soils.

## Introduction

Nitrous oxide (N_2_O) is a long-lived ozone-depleting greenhouse gas with a global warming potential that far exceeds that of the equivalent amount of CO_2_ [1, 2]. The global atmospheric N_2_O concentration has increased from 270 parts per billion (ppb) in 1750 to 331 ppb in 2018 [3]. During the 2007 to 2016 time period, the net global atmospheric N_2_O increase was estimated at 4.3 Tg N year^-1^ [4], indicating that N_2_O sources outpace N_2_O sinks. Major sources of N_2_O include denitrification (NO_3-_/NO_2-_→N_2_O) [5] and chemodenitrification [6], with additional N_2_O released from nitrification (NH_4+_→NO_3-_) [7] and dissimilatory nitrate reduction to ammonium (DNRA) (NO_3-_/NO_2-_→NH_4+_) [8]. Compared to multiple sources of N_2_O, its consumption catalyzed by N_2_O reductase (NosZ) is the only known natural biotic sink.

Canonical, complete denitrifiers possess the *nos* operon and can synthesize NosZ responsible for N_2_O reduction to dinitrogen (N_2_), the latter a gas without warming potential. Genomic analyses distinguished two types of *nos* operons with distinct *nosZ*: clade I *nosZ* generally associated with canonical denitrifying bacteria and clade II *nosZ* often found on genomes lacking the denitrification biomarker genes *nirS* or *nirK* [9–11]. Subsequent studies reported that clade II *nosZ* are generally more abundant and diverse in soils than clade I *nosZ* [12, 13], suggesting N_2_O reduction potential outside the group of complete denitrifiers. Kinetic studies using axenic bacterial cultures demonstrated that clade II N_2_O reducers exhibit higher affinities to N_2_O and growth yields than their N_2_O-reducing clade I counterparts [14], suggesting clade II reducers capture energy during growth with N_2_O as electron acceptor more efficiently. Environmental factors, including pH, temperature, O_2_ levels, substrate availability, and NO_3-_/NO_2-_ levels are known to affect N_2_O reduction [15–17]. The final reduction step (N_2_O→N_2_) catalyzed by NosZ was found particularly sensitive to pH, explaining N_2_O emissions from acidic environments [18, 19]. Expression studies suggested post-transcriptional interference at low pH, with the NosZ remaining in the catalytically inactive apo-form, a possible reason for the observed decline in NosZ activity under acidic conditions [18, 20].

Natural soils are the dominant source of N_2_O (∼5.6 Tg N_2_O-N year^-1^), accounting for approximately 33% of global N_2_O emissions [4], with tropical forest soils contributing ∼1.34 Tg N_2_O-N year^-1^ [21]. Tropical soils were reported to emit N_2_O at a rate of 0.1±0.04 g N_2_O-N m^-2^ year^-1^, which is about 50% above the average rate of global soil N_2_O emissions [22]. The large contribution of tropical soils to N_2_O emissions is explained by high N_2_ fixation activity and generally acidic soil pH [23–25]. N_2_O production can be sporadic suggesting that fluxes and concentrations of N_2_O can vary substantially both over temporal and spatial scales [26]. A comparative metagenomic study found similar relative abundances of *nosZ* sequences in acidic tropical and circumneutral temperate soils [23]. Apparently, acidic tropical soil environments have the metabolic potential to reduce N_2_O, a hypothesis supported by isotopic measurements that revealed biotic reduction of N_2_O in acidic tropical forest soils [24]. Although some evidence for N_2_O reduction under acidic conditions exists, the consensus backed by laboratory observations is that N_2_O reduction is negligible at acidic pH [18, 20, 27, 28].

To reconcile existing inconsistencies between field measurements and laboratory studies, and to explore the impacts of pH and N_2_O concentration, acidic (pH 4.4-5.0) soil samples were collected in the Luquillo Experimental Forest (LEF) in Puerto Rico. A series of microcosms explored the microbial community responses to pH (i.e., 4.5 versus circumneutral) and high versus low N_2_O concentrations. Metagenome analyses indicate that pH selects for distinct N_2_O reducers and suggest widespread distribution of N_2_O reduction potential across various acidic soil ecosystems.

## Materials and Methods

### Soil collection

LEF soil samples were collected from four locations (5-20 cm depth) in the Luquillo Experimental Forest (LEF) in Puerto Rico, including the El Verde tabonuco forest (EV, 453 m above mean sea level [MSL]), the Palm Nido palm forest (PN, 634 m MSL), the Pico del Este elfin forest (PE, 953 m MSL), and the Sabana tabonuco forest (S, 265 m MSL) (Fig. S1). The soil materials were transferred to sterile Whirl-Pak bags, placed at 4°C, and manually homogenized prior to microcosm setup. Detailed descriptions of the LEF can be found elsewhere [29, 30], and physicochemical properties of the soil samples are presented in Table S1.

### Soil microcosms

Completely synthetic, reduced (0.2 mM L-cysteine) mineral salt medium was prepared in 160 mL glass serum bottles following established protocols [31]. In the pH 4.5 medium, 50 mM potassium dihydrogen phosphate replaced the 30 mM bicarbonate buffer used in the pH 7.3 medium [32]. The pH was adjusted with CO_2_ (pH 7.3 medium) or with 4 M hydrochloric acid (pH 4.5 medium). CuCl_2_ (17 μM) and the Wolin vitamin mix were added from concentrated stock solutions to individual vessels after autoclaving [32].

Inside a glove box (Coy Laboratory Products, Grass Lake, MI, USA) filled with 97% N_2_ and 3% H_2_, 2 g (wet weight) of homogenized soil material was aseptically transferred to glass serum bottles containing 100 mL of medium using sterilized stainless-steel spatulas. The serum bottles were immediately resealed with sterile butyl rubber stoppers, crimped with aluminum caps, and removed from the glove box. Lactate (5 mM) was added to each serum bottle from 1 M stock by syringe and replenished four times over the 15-month incubation period according. Two replicate series of microcosms were established at pH 4.5 and at pH 7.3 with duplicate microcosms maintained for the pH 4.5 incubations (Table 1 and Fig. S1). Plastic syringes (BD, Franklin Lakes, NJ, USA) with needles (25-gauge BD, Franklin Lakes, NJ, USA) were used to add 0.1 mL (4.17 µmol, 0.02 mM aqueous N_2_O) and 10 mL (416.7 µmol, 2 mM aqueous N_2_O) of undiluted N_2_O gas to the incubation vessels. N_2_O was periodically analyzed by gas chromatography (GC) as described [32] and replenished when consumed. All bottles were incubated at 30°C under static conditions for 15 months. Negative controls included heat-killed (autoclaved) replicates and microcosms without N_2_O but with lactate at pH 4.5 for each soil sample.

### Analytical procedures

N_2_O was analyzed with an Agilent 3000A Micro-GC (Agilent, CA, USA) equipped with a thermal conductivity detector and a Plot Q column [14]. The limit of detection was 50 ppmv N_2_O with signal-to-noise ratio of 3:1. The injector and column temperatures were set to 100°C and 50°C, respectively, and the column pressure was set to 25 psi. For each measurement, a 0.1 mL headspace sample was withdrawn from the microcosm vessel and manually injected into the Micro-GC. Aqueous N_2_O concentrations were calculated from the headspace concentration using a dimensionless Henry’s constant for N_2_O at 20°C of 1.68 based on the equation *C*_aq_ = *C*_g_/*H*_cc_ [33]. *C*_aq_ and *C*_g_ are the aqueous N_2_O and the headspace N_2_O concentrations (μM), respectively, and *H*_cc_ is the dimensionless Henry’s constant. The total amount of N_2_O was calculated as the sum of N_2_O in headspace and aqueous phases.

### DNA extraction and metagenome sequencing

When about half of the final N_2_O amendment had been consumed, the microcosms were shaken, and 5 mL suspension samples were collected with 5-mL plastic syringes equipped with 18-gauge needles. DNA for shotgun metagenome sequencing was extracted with the DNeasy PowerSoil kit (Qiagen, Hilden, Germany) according to the manufacturer’s instructions. DNA concentrations were determined using the Qubit fluorometer (Life Technologies, Carlsbad, CA, USA). Sequencing of the metagenomic DNA was performed at the Institute for Genome Sciences at the University of Maryland using the Novaseq 6000 platform (Illumina, San Diego, CA, USA) to generate 48 to 73 million reads with 150-bp read length per sample (Table S2).

### Bioinformatic analysis

The metagenomes of the four original soils were sequenced previously [23], and were downloaded from European Nucleotide Archive under project PRJEB26500. The original soil raw reads and the 16 N_2_O-reducing microcosms were trimmed using Trimmomatic v0.39 using default parameters [34]. Subsequent assembly was performed using IDBA-UD v1.1.3 [35], and only contigs longer than 1,000 bp were included in downstream analyses. Contigs were binned using MaxBin2 v2.2.4 with default settings to recover individual MAGs [36]. MAGs were dereplicated with dRep using default parameters [37] and checked for completeness and contamination using CheckM v1.0.18 [38], before mapping was performed with Bowtie2 using default settings [39]. The resulting MAGs were evaluated for their intrapopulation diversity and sequence discreteness using fragment recruitment analysis scripts available through the Enveomics collection [40].

### Metagenomic community profiling

GraftM v0.13.1 was used to extract 16S rRNA gene fragments from the trimmed metagenomic datasets for classification using the Greengenes database (release 13_8) at the 97% nucleotide identity level [41, 42]. The relative abundance of the operational taxonomic units (OTUs) was calculated based on the number of reads assigned to each OTU. Community profiling was based on OTU taxonomic assignments at the phylum, family, and genus levels.

### Identification of *nosZ* genes

ROCker was used to identify metagenomic reads carrying *nosZ* [43]. Briefly, trimmed short reads were used as the query for BLASTX (Diamond v0.9.14.115) searches against the corresponding ROCker protein database representing the target gene [44]. The matching sequences were then filtered using the ROCker compiled models available at http://enve-omics.ce.gatech.edu/. The abundances of target genes (i.e., clade I and clade II *nosZ*) were determined by calculating the ratio between normalized target reads (counts divided by the median protein length) and the genome equivalents [45]. The genome equivalents of each metagenomic dataset were obtained using the MicrobeCensus package [46]. Reference *nosZ* genes were also searched against the assemblies and MAGs using precompiled hidden Markov models obtained from FunGene and HMMer [47, 48]. Hits with an identity value of 100% were filtered based on the NosZ sequences in the reference database [23].

### NosZ phylogeny

NosZ reference sequences were aligned with ClustaloΩ using default settings [49]. The alignment was used to build a maximum likelihood reference tree in RAxML V8.2.12 with ‘-f a’ algorithm, gamma parameter optimization, and a general time reversible model option [50]. ROCker identified reads carrying *nosZ*, which were translated to protein sequences using MetaGeneMark [51]. The translated sequences were added to the NosZ reference protein alignment using MAFFT and the ‘addfragments’ option [52], and the new alignment placed in the NosZ reference phylogenetic tree using RAxML EPA algorithm (-f v option). The generated jplace file was processed using an in house script (available through http://enve-omics.ce.gatech.edu/) for visualization in iTOL [53]. Detected NosZ proteins in the assemblies and MAGs were manually curated, added to the phylogenetic trees, and visualized as described above.

### Taxonomic assignment, functional annotation, and comparative genomics

Taxonomic assignments of MAGs with >75% completeness and <5% contamination used the GTDB-Tk v0.1.4 tool [54] of the Genome Taxonomy Database (GTDB, http://gtdb.ecogenomic.org) version R202 [55]. Protein-coding sequences were predicted using Prodigal v2.6.3 [56], and assigned KEGG orthologs using KofamScan v1.3.0 against HMM profiles from the KEGG database (released on 24-Feb-2021) [57]. The completeness of various metabolic pathways was assessed using KEGG-Decoder v1.32.0 [58], and pathways of interest (e.g., nitrogen cycling, lactate and hydrogen metabolism, copper transport) were manually selected from the KEGG-Decoder results. Average nucleotide identity (ANI) and average amino acid identity (AAI) between high-quality MAGs and genomes of phylogenetically related bacteria were calculated with MiGA (http://microbial-genomes.org/). Phylogenetic trees were created using FastTree 2.1.8 (WAG+GAMMA models) with a concatenated alignment of 120 bacterial and 122 archaeal conserved marker genes [59] and visualized with iTOL [53].

### N_2_O reduction potential in acidic soils

From MAGs and contigs of acidic N_2_O-reducing microcosms, 33 near full-length NosZ sequences were recovered (Fig. S2 and Fig. S3), and dereplication yielded eight unique NosZ sequences. A customized database comprising 14 NosZ included six closely related sequences from the reference NosZ database. To identify the respective *nosZ* genes in various acid (pH 3.5-7.5) soil microbiomes, 35 available soil metagenome datasets were downloaded from the European Nucleotide Archive (Table S3) and subjected to BLASTX query using the customized NosZ database with a minimum identity value of 60% and an e-value of 1e-05. The normalization approach described above was used to calculate relative abundances based on *nosZ*-carrying metagenomic reads.

### Statistics

Statistical analyses were performed using R version 4.0.2. Beta-diversity was calculated using Bray–Curtis dissimilarity and visualized using the principal coordinate analysis (PCoA) plot in R with packages ggplot2 [60] and phyloseq [61]. Statistical differences in microbial communities among original soils, pH 4.5 microcosms, and pH 7.3 microcosms were determined using permutational multivariate analysis of variance (PERMANOVA) using the adonis function in vegan with 999 permutations [62]. Heatmaps of annotation results of MAGs were generated using the R package pheatmap.

## Results

### N_2_O reduction in tropical forest soil microcosms

N_2_O consumption occurred with lag times of 1-2 weeks in pH 4.5 and pH 7.3 microcosms with low level of N_2_O (0.02 mM aqueous concentration). Longer lag times of up to 7 months were observed in microcosms with 2 mM aqueous N_2_O under both pH conditions, but microcosms with high-level N_2_O consumed substantially more N_2_O over the 15-month incubation period (Fig. S4 and Table S4). No N_2_O loss occurred in autoclaved microcosms, indicating the acidic tropical forest soils harbor N_2_O-reducing microorganisms. In addition, no N_2_O was detected in control microcosms that did not receive N_2_O, indicating that N_2_O formation from nitrogenous compounds in the medium or associated with the soil did not occur or was negligible.

### Bacterial community composition

Following the 15-month incubation period and when the microcosms had consumed about half of the final N_2_O feeding, DNA was extracted for metagenome sequencing. Totals of 5,963 ± 2,503 and 23,399 ± 15,030 sequences representing 16S rRNA genes were obtained from the original soil and the corresponding microcosm metagenome datasets, respectively, and yielded 903 16S rRNA gene-based OTUs (Table S5).

Rarefaction analysis suggested that the number of unique 16S rRNA genes approached saturation in most samples (Fig. S5), and the number of OTUs detected in N_2_O-reducing microcosms was lower than in the corresponding original soils. Beta diversity analysis using the 16 metagenomic datasets generated from the microcosms as well as the four datasets generated from the original soils [23] indicated distinct community compositions in response to pH and N_2_O levels (*p* < 0.01, PERMANOVA) (Fig. 1A, Table S6). PCoA revealed that ∼55% of the total variability of OTUs observed in pH 4.5 and in circumneutral N_2_O-reducing microcosms compared to the respective original soils was explained by pH and N_2_O (Fig. 1A). Datasets acquired from same pH microcosms established with the four different soils clustered together indicating that pH shaped distinct microbial communities over the 15-month incubation period (Fig. 1).

**Fig. 1.**
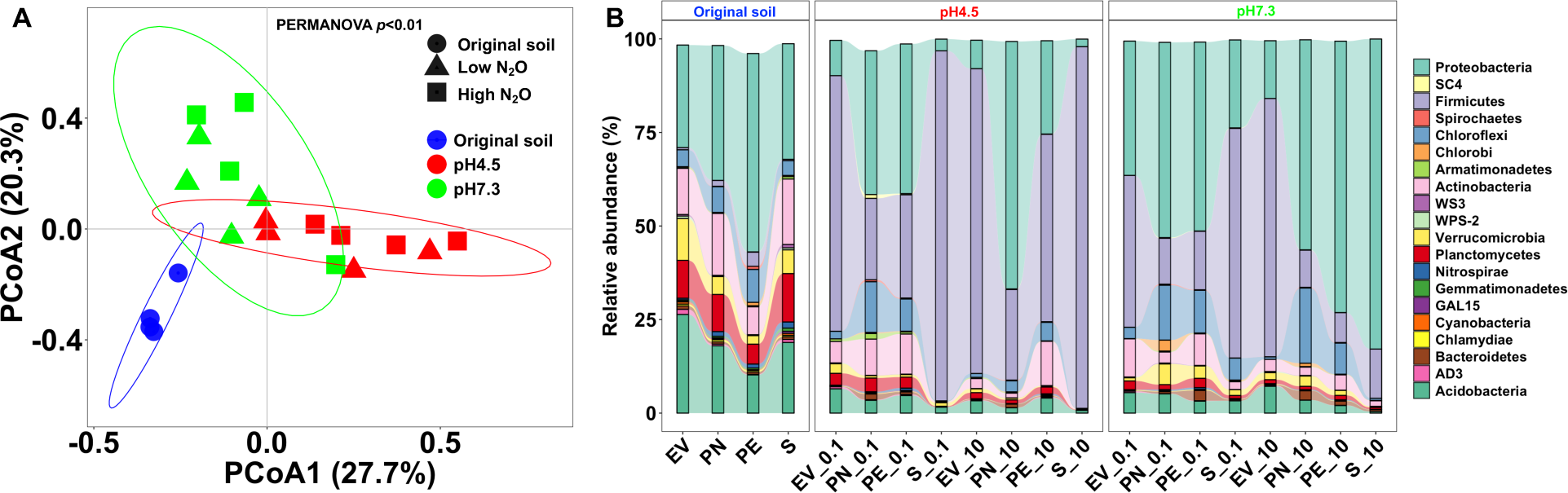
Microbial community composition of the original tropical forest soils and the N_2_O-reducing microcosms maintained under low versus high levels of N_2_O and at pH 4.5 versus pH 7.3. (A) Beta diversity of microbial communities based on weighted Unifrac analysis of 16S rRNA gene fragments recovered from the metagenomes. Samples are visualized by PCoA with colors distinguishing original soils and microcosms at acidic versus circumneutral pH. The ellipses represent the 95% confidence intervals. (B) The relative abundance distributions of the top 20 phyla observed in the original soils and the different microcosms. The numerals shown on the x-axis labels indicate the volumes of N_2_O added with each feeding to the microcosms (0.1 mL [0.02 mM aqueous N_2_O] versus 10 mL [2 mM aqueous N_2_O]).

The majority of the 16S rRNA gene sequences derived from the original soils affiliated with the phyla *Proteobacteria*, *Actinobacteria*, *Verrucomicrobia*, *Planctomycetes*, and *Acidobacteria*, with a combined relative abundance exceeding 70% (Fig. 1B). Higher relative abundances of *Proteobacteria*, *Firmicutes*, *Chloroflexi*, and *Actinobacteria* were observed in the 16 N_2_O-reducing microcosms relative to the original soil inoculum. Analysis at the family level showed that sequences representing *Peptococcaceae*, *Veillonellaceae*, *Clostridiaceae*, and *Hyphomicrobiaceae* increased in acidic microcosms, and sequences of *Rhodocyclaceae*, *Clostridiaceae*, *Hyphomicrobiaceae*, and *Ruminococcaceae* were more abundant in microcosms maintained at circumneutral pH compared to the original soils (Fig. S6A). Based on the relative abundances of 16S rRNA gene sequences (highest observed values shown in parentheses), the genera *Desulfosporosinus* (45%), *Desulfomonile* (5%), *Rhodoplanes* (53%), *Azospira* (56%), and *Dechloromonas* (38%) increased in response to N_2_O additions compared to the original soils (Fig. S6B). At least some members of these genera comprise known N_2_O-respiring species [9], suggesting that the bacteria capable of using N_2_O as an electron acceptor were enriched.

### Phylogenetic distribution and relative abundance of clade II versus clade I *nosZ*

N_2_O reduction is catalyzed by NosZ and enrichment with N_2_O increased the *nosZ* gene abundances. The analysis of the metagenome datasets showed that clade II *nosZ* sequences outnumbered clade I *nosZ* gene sequences in the original soils and in the N_2_O-reducing microcosms (except the S_pH7.3_0.1) (Fig. S7, Table S7). Placing the extracted clade II *nosZ* sequence reads from the original soils in the reference clade II *nosZ* phylogenetic tree revealed that most of the reads extracted from the original soil metagenome datasets affiliated with the genera *Anaeromyxobacter* and *Opitutus* (Fig. 2A). In contrast, the *nosZ* sequence reads observed in the acidic microcosms were assigned to the genera *Profundibacter*, *Desulfosporosinus*, and *Desulfomonile* (Fig. 2B and C). In microcosms maintained at pH 7.3, the majority of *nosZ* reads affiliated with the genera *Azospira*, *Dechloromonas*, and *Sulfuricella* (Fig. 2D and E), except for the EV soil microcosms with high level of N_2_O, where *nosZ* sequences assigned to the genus *Desulfosporosinus* dominated. A comparative analysis of clade II *nosZ* reads revealed that the microcosms maintained at pH 4.5 and at pH 7.3 developed distinct N_2_O-reducing communities (*p* < 0.01, PERMANOVA) (Fig. S8A). Taken together, these analyses suggest that bacteria with clade II *nosZ* drive N_2_O reduction in all tropical soil microcosms, but pH selects for distinct clade II N_2_O reducers (Fig. 2).

**Fig. 2.**
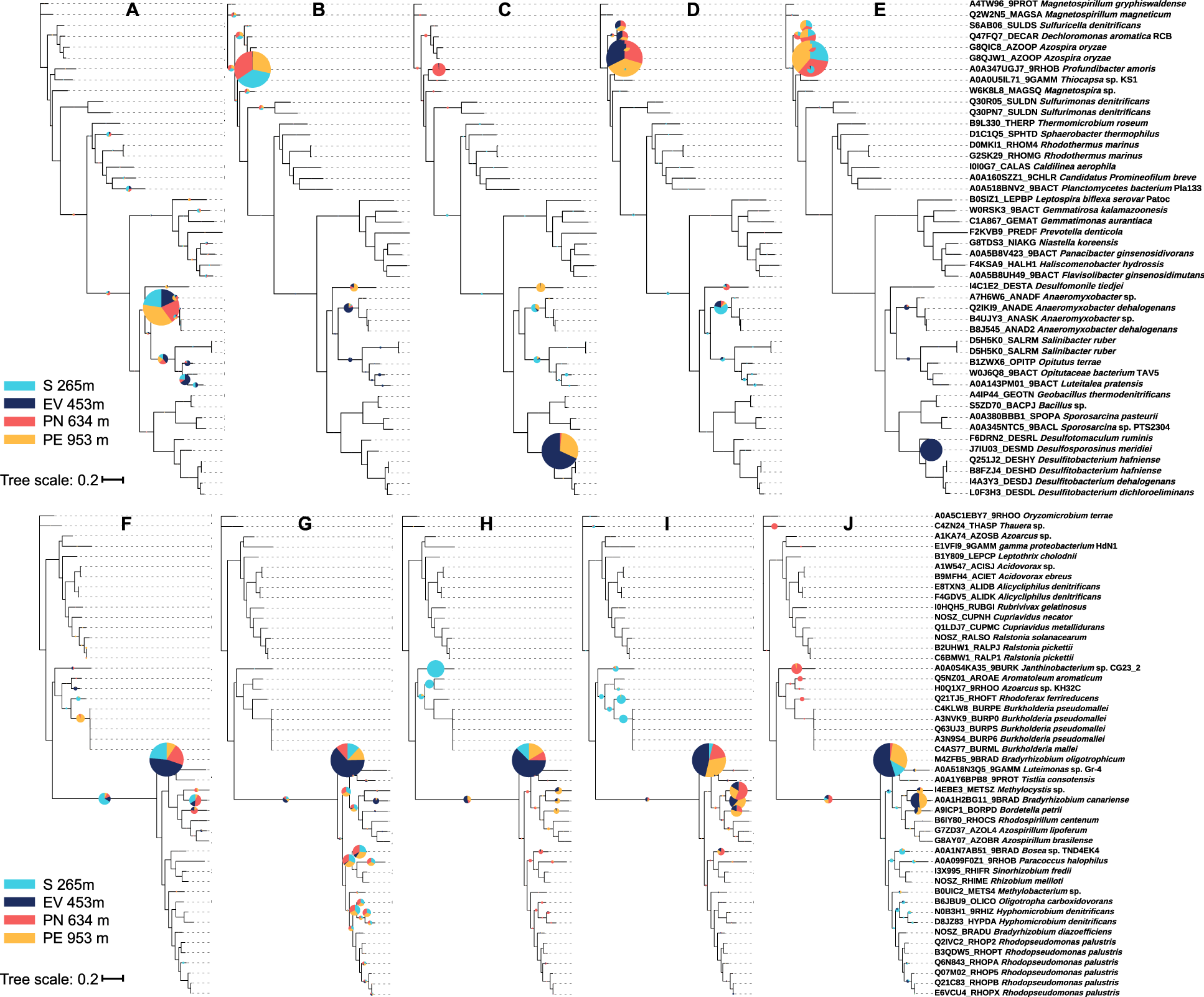
Phylogenetic diversity of *nosZ* reads recovered from the four original soils and the 16 microcosms maintained at pH 4.5 versus pH 7.3 and at low versus high levels of N_2_O. Trimmed clade II and clade I *nosZ* reads in the metagenomes were identified by ROCker and placed in the corresponding reference *nosZ* phylogenetic tree, as described in the Methods section. The radii of the pie charts are proportional to the number of reads assigned to each *nosZ* subclade and the colors represent the different soils. (A) Diversity of clade II *nosZ* reads from metagenomes of the four original soil samples. (B) Phylogenetic information of clade II *nosZ* in N_2_O-reducing microcosms at acidic pH with low level of N_2_O. Phylogenetic information of clade II *nosZ* in microcosms at acidic pH with high level of N_2_O (C), at circumneutral pH with low level N_2_O (D), and at circumneutral pH with high level N_2_O (E). (F) Diversity of clade I *nosZ* reads from metagenomes of the four original soil samples. (G) Phylogenetic information of clade I *nosZ* in N_2_O-reducing microcosms at acidic pH with low level of N_2_O. Phylogenetic information of clade I *nosZ* in microcosms at acidic pH with high level of N_2_O (H), at circumneutral pH with low level N_2_O (I), and at circumneutral pH with high level N_2_O (J).

Most of the clade I *nosZ* sequences could be assigned to the genus *Bradyrhizobium*, and a small number of reads affiliated with the genera *Methylocystis*, *Methylocella*, and *Janthinobacterium*, independent of the pH conditions or N_2_O levels (Fig. 2F, G, H, I, and J). Apparently, acidic versus circumneutral pH and low versus high N_2_O levels did not select for distinct clade I *nosZ* N_2_O reducers. The PCoA supports that the N_2_O-reducing communities (clade I) in acidic pH and circumneutral pH microcosms were similar (*p* > 0.05, PERMANOVA) (Fig. S8B).

### Metagenome-assembled genomes (MAGs)

A non-redundant set of 17 high-quality MAGs (>75% completeness and <5% contamination) harboring *nosZ* genes was recovered from the 16 metagenome data sets generated from the N_2_O-reducing microcosms (Fig. 3A, Tables S8 and S9). By mapping the trimmed metagenomic reads from each N_2_O-reducing microcosm to the MAGs harboring *nosZ* genes, between 1.3% and 44% of the sequence reads mapped to the corresponding MAGs, revealing that these MAGs represented abundant members of the enriched community (Fig. 3B). Only five MAGs with *nosZ* were detected in the metagenomic data of the original soils with a relative abundance of 0.01-0.02% (Fig. S9), indicating N_2_O-reducing microbes were strongly enriched in microcosms that received N_2_O.

**Fig. 3.**
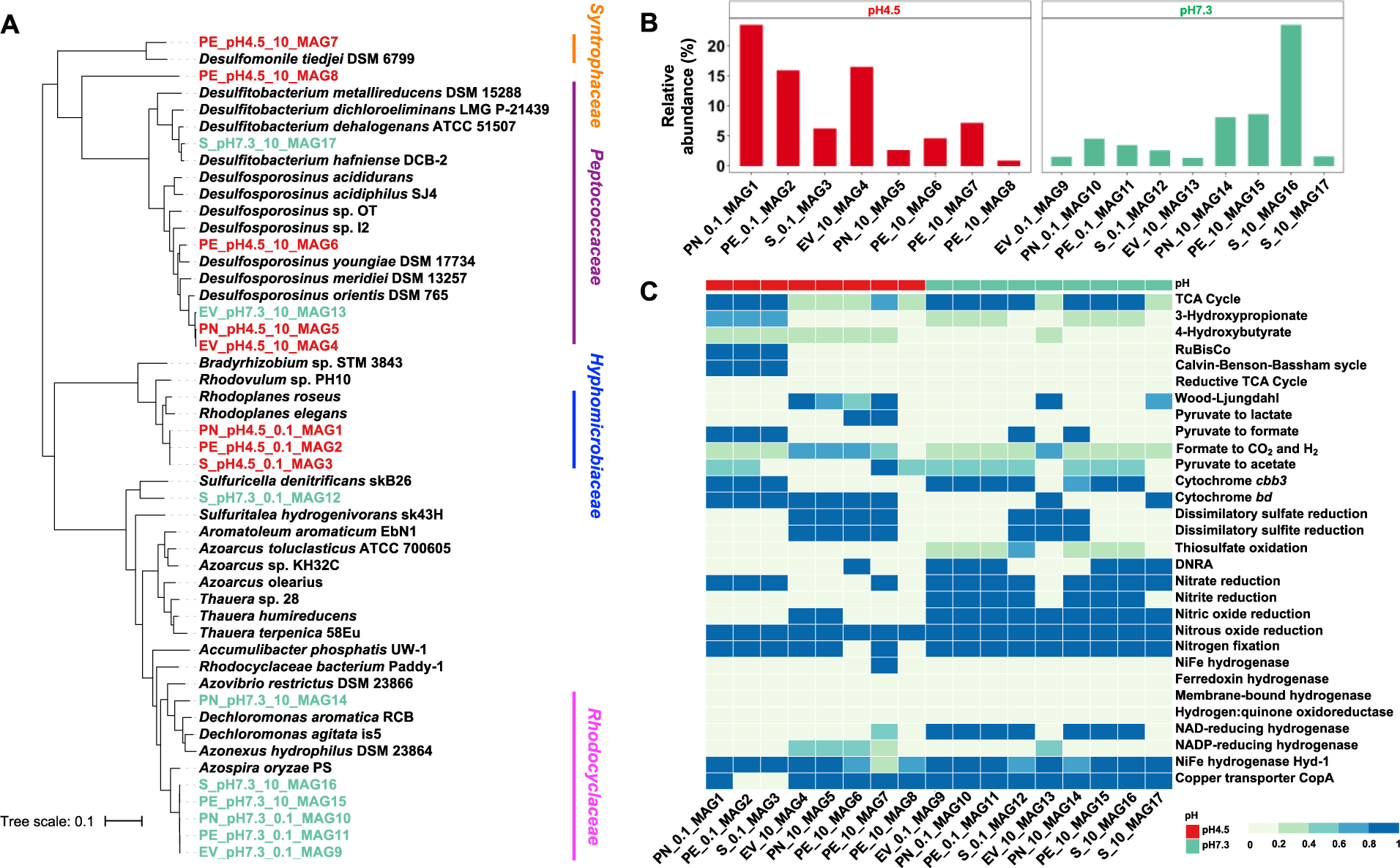
Phylogenetic and functional analysis of MAGs harboring *nosZ* genes. Panel (A) shows a phylogenetic tree of the 17 MAGs harboring a *nosZ* gene and their closest neighbors based on the analysis of 120 bacterial marker genes. MAGs recovered from pH 4.5 and pH 7.3 N_2_O-reducing microcosms are shown in red versus green font, respectively. Panel (B) shows the relative abundance of each MAG harboring a *nosZ* gene in the corresponding N_2_O-reducing microcosms based on metagenomic reads competitively mapped against the MAG sequences. The abundance of these MAGs in the original soil metagenomes was below 0.01% (Fig. S9). Panel (C) depicts a heatmap showing the completeness of key metabolic pathways or functions in the 17 MAGs harboring a *nosZ* gene based on KEGG annotation. The numbers 0.1 and 10 in the x-axis labels represent the low and high levels of N_2_O added to the microcosms, respectively.

All 17 MAGs harboring *nosZ* genes were assigned to novel taxa according to the AAI results (Table S10), indicating an unexplored diversity of N_2_O reducers in the tropical forest soils and enriched in the N_2_O-reducing microcosms. Six of eight MAGs recovered from acidic N_2_O-reducing microcosms were taxonomically close to the genera *Desulfitobacterium*, *Rhodoplanes*, *and Desulfosporosinus* with AAI values exceeding 68% (Table S10), probably representing novel species [63]. The AAI similarities between the remaining two MAGs, PE_pH4.5_10_MAG7 and MAG8, and the corresponding closest relatives are 45.04 % and 44.04 %, respectively, indicating these MAGs most likely represent novel families (Fig. 3A and Table S10). MAGs obtained from circumneutral, N_2_O-reducing microcosms were taxonomically related to the genera *Azospira*, *Sulfurimicrobium*, *Dechloromonas*, and *Desulfitobacterium* (Table S10).

### Functional analysis of MAGs harboring *nosZ* genes

Key metabolic pathways or functions of the 17 MAGs harboring *nosZ* were predicted based on the KEGG annotation (Fig. 3C). All eight MAGs obtained from acidic N_2_O-reducing microcosms lack the hallmark denitrification genes *nirK*/*nirS* encoding nitrite reductase, indicating these organisms are non-denitrifiers; however, seven of the nine MAGs derived from circumneutral microcosms carried a complete set of genes for canonical denitrification (i.e., NO_3-_→NO_2-_→N_2_O→N_2_). All MAGs representing complete denitrifiers harbored *nirS*. In addition to *nosZ*, other genes of the *nos* operon were detected in all MAGs (Fig. S10). These findings indicate that non-denitrifying N_2_O-reducing bacteria may be the main drivers for N_2_O reduction in low pH soils. NosZ is a copper-containing enzyme and the extracellular copper concentration controls *nosZ* expression [64]. Notably, genes for copper transport were identified in 14 of the 17 MAGs harboring *nosZ*.

Sources of electrons for reductive processes (i.e., N_2_O reduction) were exogenously added lactate and organic material associated with the soil. Lactate was readily consumed in all live microcosms. The analysis of two MAGs (PE_pH4.5_10_MAG6 and MAG7) recovered from acidic N_2_O-reducing microcosms revealed genes encoding L-lactate dehydrogenase (*LDH*, K00016) implicated in the conversion of lactate to pyruvate, but only PE_pH4.5_10_MAG6 contained the complete set of genes (*pta*, K13788, and *acyP*, K01512) required to metabolize lactate to acetate (Fig. 3C). In addition, five MAGs, three from acidic and two from circumneutral pH microcosms, possessed genes implicated in the fermentation of pyruvate to formate and acetyl-CoA (Fig. 3C). These findings suggest that most MAGs harboring *nosZ* genes were unable to directly utilize lactate as an electron donor. Collectively, these results demonstrate that MAGs harboring *nosZ* genes are taxonomically and metabolically diverse.

### *nosZ* genes in metagenomes representing low pH soil biomes

A total of 27 metagenomes representing low pH (pH 3.5-5.7) forest, agricultural, and permafrost soil ecosystems were analyzed (Table S3), and all of them harbored *nosZ* sequences. *nosZ* abundances ranged from 0.002 ±0.001 to 0.25 ±0.11 genome equivalents (i.e., the fraction of genomes expected to carry *nosZ* assuming one gene copy per genome; Fig. 4), suggesting that N_2_O reduction potential is common in low pH soils. The highest abundances of *nosZ* genes were observed in permafrost soils, and the lowest abundances were detected in two temperate forest soils. *nosZ* genes were also abundant in low pH tropical forest soils, which typically have high N_2_ fixation and nitrogen turnover activities. An expanded survey that included the 27 metagenomic datasets from low pH plus eight datasets from pH 6–7.5 soils revealed that *nosZ* sequences representing low pH soil biomes were predominantly found in pH 4.5–6 environments (Fig. S11). The observation that *nosZ* genes commonly occur in acidic soils suggest that the potential for low pH microbial N_2_O reduction is broadly distributed.

**Fig. 4.**
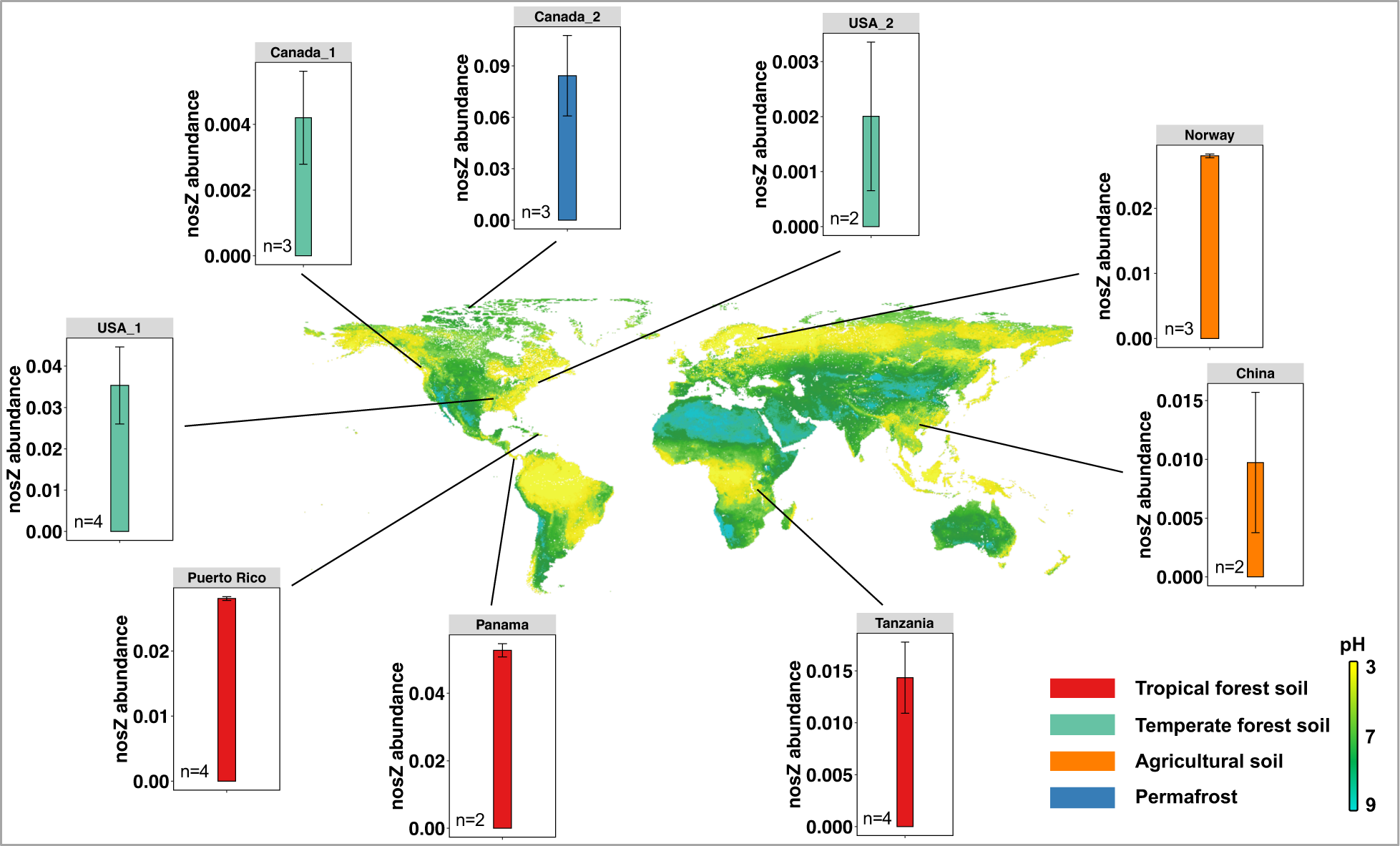
Distribution of N_2_O reduction potential in acidic (pH 3.5-5.7) soils. Detailed information about the metagenome datasets is provided in Table S3. Global soil pH data were obtained from the Soil Geographic Databases (https://www.isric.org). The plots show the total abundance of *nosZ* genes per genome equivalent. The black lines point to the approximate locations from where the soil metagenomes were derived.

## Discussion

### pH paradigm for microbial N_2_O reduction

Biological N_2_O reduction is catalyzed by microorganisms expressing NosZ with maximum efficiency observed at circumneutral pH [65, 66]. N_2_O reduction under acidic pH conditions has received considerable attention; however, experimental work with laboratory cultures indicated that NosZ activity seizes at pH <5.7 [18, 20, 27, 28]. These observations have been used to explain the lack of N_2_O reduction activity under low pH conditions, and acidic environments are generally considered N_2_O sources. NosZ is a periplasmic enzyme, and biochemical studies suggested that low pH impacts the assembly and maturation of functional NosZ [18], a plausible explanation for the observed lack of N_2_O reduction activity at acidic pH. In contrast to the observations with laboratory cultures, soil microcosms studies reported N_2_O reduction at pH <5.7 [67–69]. A possible reason for the contradictory findings is the presence of microsites within the soil matrix with higher pH than the measured bulk aqueous phase pH, allowing microbial N_2_O reduction to occur in such micro- environments [18, 70, 71]. Further, the interpretation of observations made in short-term soil microcosm incubations is not straightforward because it is unclear if the N_2_O reduction activity observed under acidic pH conditions is due to residual activity of existing biomass, or linked to the formation of new cells that conserve energy for growth from N_2_O reduction. The sustainable reduction of N_2_O requires the formation of new biomass (i.e., growth) under the prevailing environmental conditions. Laboratory studies with consortia that utilize the toxin vinyl chloride as respiratory electron acceptor illustrate this issue. Reductive dechlorination activity was observed at pH <5.5 [70]; however, this activity relied on existing biomass produced at circumneutral pH, and growth of vinyl chloride-respiring *Dehalococcoides mccartyi* strains did not occur at acidic pH, indicating that a sustainable process in acidic groundwater cannot be envisioned [70]. In analogy, it is uncertain if the results of short-term microcosm studies without repeated N_2_O feedings generate meaningful information to predict *in situ* N_2_O reduction activity in low pH soils. Further complicating data interpretation is the observation that common oxygen-respiring bacteria (e.g., members of the *Gemmatimonadaceae*) utilize N_2_O as an electron sink following oxygen depletion; however, this process is uncoupled from growth and not sustained under anoxia [72, 73]. The LEF soil microcosms were incubated for 15-months and repeated N_2_O additions were consumed at accelerating rates, an observation inconsistent with activity of residual (i.e., non-growing) biomass, and indicative of respiratory N_2_O utilization and growth of N_2_O reducers at pH 4.5. Our observations challenge the notion that efficient N_2_O reduction requires circumneutral pH and suggest that N_2_O reduction can be sustained under acidic pH conditions.

### pH selects for distinct N_2_O reducers

Prior work demonstrated that LEF soils collected at different locations share the predominant clade II *nosZ* genes and implicated similar taxa in N_2_O reduction [23]; however, the microcosm study revealed differences in the taxa responsible for N_2_O reduction (i.e., clade II type N_2_O reducers) depending on the elevation where the soil samples were collected (Figs. 2 and 3). For instance, the metagenome analysis of the original soils prior to enrichment implicated *Anaeromyxobacter* populations as predominant N_2_O reducers (Fig. 2). N_2_O reduction by *Anaeromyxobacter* at circumneutral pH has been demonstrated [6]; however, the predominant N_2_O reducers in microcosms maintained at circumneutral pH selected for different N_2_O-reducing taxa, including *Azospira* and *Dechloromonas* species, which have reported growth optima near pH 7.0 [74, 75]. Apparently, circumneutral pH favored *Azospira* and *Dechloromonas* over N_2_O-reducing *Anaeromyxobacter* species in the microcosms, possibly due to kinetic differences. The relative abundance of *Anaeromyxobacter* sequences in the original soils can be explained by the presence of ferric iron, a favorable electron acceptor for *Anaeromyxobacter* [6]. In acidic N_2_O-reducing microcosms, sequences representing the genera *Desulfosporosinus*, *Desulfitobacterium*, *Desulfomonile*, and *Rhodoplanes* increased. These genera comprise members that grow under acidic conditions (e.g., pH 4-6), and phenomic studies have shown that at least some affiliated taxa can reduce N_2_O [76–79]. The tropical soil microcosm experiments demonstrate that pH selects for different N_2_O-reducing taxa harboring clade II *nosZ*, and show that N_2_O reducers are distributed along the LEF elevational gradient spanning ∼700 m. In addition, specific N_2_O reducers enriched in the microcosms were rare in the four original soils, an observation not uncommon in soil enrichment studies and possible indicator for a diverse community of active N_2_O reducers in situ. Despite the presence of taxa capable of low pH N_2_O reduction, acidic tropical forest soils are considered N_2_O emitters, raising the question of parameters, other than pH, that are limiting N_2_O reduction activity in this relevant terrestrial ecosystem.

### Non-denitrifying N_2_O reducers responsible for N_2_O reduction under low pH conditions

A recent metagenome analysis of the same LEF soils used for microcosm setup found abundant *nosZ* genes, with clade II *nosZ* genes generally more abundant than clade I *nosZ* genes [23].

Following the 15-month incubation period, clade II *nosZ* dominated the *nosZ* gene pools in all microcosms irrespective of pH and N_2_O concentration (Fig. 2 and Fig. S7), indicating that the enrichment conditions favored populations with clade II *nosZ*. The analysis of MAGs derived from acidic N_2_O-reducing microcosms harboring *nosZ* indicated that non-denitrifiers (i.e., bacteria lacking *nirS*/*nirK*) that represent novel taxa (Table S10) are responsible for N_2_O reduction (Fig. 3 and Fig. S3). Prior studies reported that N_2_O reduction under slightly acidic conditions (pH ∼6.0) was driven by complete denitrifiers that represent cultivated taxa [80, 81]. Consistent with these prior reports, the MAGs with *nosZ* derived from microcosms maintained at circumneutral pH represent complete denitrifiers. Of note, three MAGs derived from acidic microcosms with low level of N_2_O each harbored two *nosZ* genes, one clade I and one clade II, a genotype observed for few species of the class *Betaproteobacteria* capable of denitrification at circumneutral pH [82]. Collectively, the findings show that circumneutral pH selects for complete denitrifiers, whereas acidic pH selects for non-denitrifying N_2_O reducers in the tropical forest soils studied here. A relevant conclusion from this observation is that complete denitrifiers do not represent good models to study low pH N_2_O reduction.

### Impact of N_2_O concentration on N_2_O reduction

A striking difference in microcosms with 0.02 mM versus 2 mM N_2_O was the extended lag phase observed in the high-level N_2_O microcosms independent of the pH condition. A possible explanation for the delayed start of N_2_O consumption is the inhibition of corrinoid-dependent pathways. Micromolar concentrations of N_2_O were shown to repress methionine biosynthesis [64], methanogenesis [83], methylmercury formation [84], and bacterial reductive dechlorination [32], all processes that involve enzymatic steps that strictly depend on the cobalt (I) supernucleophile, a species highly susceptible to oxidation by N_2_O [85]. An initial N_2_O concentration of 2 mM exceeds the reported inhibitory constants for corrinoid-dependent enzymes about 100-fold, suggesting the disruption of metabolic pathways directly or indirectly impacted N_2_O-consuming populations. Only a subset of bacteria and archaea synthesize corrinoids [86], and corrinoid prototrophs supply this essential nutrient to corrinoid auxotrophs [87, 88]. The addition of N_2_O disrupts these interactions, an effect influenced by the concentration of N_2_O. The delayed onset of N_2_O reduction in microcosms with high level of N_2_O may reflect a switch to corrinoid-independent pathways [32, 64].

### Implications for N_2_O reduction in low pH environments

Global atmospheric N_2_O concentrations are on the rise, and the acidification of agricultural and forest soils, two major sources of atmospheric N_2_O, is predicted to exacerbate emissions [4, 89, 90]. Soils with a pH below 5.5 currently comprise approximately 30% of the global ice-free land area and are mainly distributed in the northern temperate to cold belt and the southern tropical belt [91]. Our findings suggest that non-denitrifying microbes catalyze N_2_O reduction at acidic pH, and that microbial N_2_O reduction potential is widely distributed in acidic soils (Fig. 4). These observations seem at odds with the dogma that acidic environmental systems are predominantly N_2_O sources, and future research should address two major knowledge gaps. While N_2_O pools (i.e., how much is there?) can be quantitatively captured, robust tools to measure N_2_O fluxes (i.e., the rates of N_2_O formation and consumption) that ultimately determine emissions are lacking. Thus, it is not obvious if N_2_O emissions from acidic soil ecosystems reflect a lack of microbial N_2_O consumption capacity or an imbalance between N_2_O formation versus consumption. Very limited information is available about the taxonomic diversity, physiology, and ecology of microbes that reduce N_2_O under acidic conditions. This study identified several genera (i.e., *Desulfosporosinus*, *Desulfitobacterium*, *Desulfomonile*, and *Rhodoplanes*) as potential targets for more detailed exploration of low pH N_2_O reduction, and future efforts should focus on the isolation and characterization of N_2_O reducers from low pH environments to generate detailed process understanding.

## Acknowledgments

The authors acknowledge funding through the Dimensions of Biodiversity program of the US National Science Foundation (awards 1831582 to K.T.K. and 1831599 to F.E.L.). Y.S., Y.Y., and G.H. acknowledge the support from the China Scholarship Council.

