## Supplemental Material for "pH selects for distinct N_2_O-reducing microbiomes in tropical soil microcosms"

**Supporting Information summary:** 10 tables, 11 figures, and references.

**Table S1.** Physicochemical properties of soil samples collected along an elevational gradient in the Luquillo Experimental Forest (LEF) in Puerto Rico [1].

| Sampling location | Abbreviation | Moisture | Class | pH | Total C | Total N | $NO_3$-N | $NO_2$-N | $NH_4$-N | Sulfate |
|---|---|---|---|---|---|---|---|---|---|---|
| | | | | | % [a] | | mg kg$^{-1}$ [b] | | | |
| Sabana | S | 30.70% | Clay | 4.72 | 1.717 | 0.167 | < 0.190 | < 0.210 | 2.39 | 3.43 |
| El Verde | EV | 33.80% | Clay | 4.45 | 3.698 | 0.246 | < 0.190 | < 0.210 | 3.78 | 2.31 |
| Palm Nido | PN | 48.90% | Clay Loam | 4.97 | 6.604 | 0.280 | < 0.190 | < 0.210 | 5.52 | 10.08 |
| Pico del Este | PE | 54.20% | Silty Clay Loam | 4.67 | 8.489 | 0.338 | < 0.190 | < 0.210 | 1.16 | 6.46 |

[a] Reported as percent of dry weight soil.

[b] Reported in mg kg$^{-1}$ (i.e., wet weight of soil at the time of sampling).

S, Sabana (265 m above mean sea level [MSL]); EV, El Verde (453 m MSL); PN, Palm Nido (634 m MSL); PN, Pico del Este (953 m MSL).

Additional information about of the vegetation and soil characteristics of the LEF sampling locations are available in the published literature [2, 3].

**Table S2.** Information about the metagenome data generated from the 16 $N_2O$-reducing tropical soil microcosms.

| | | | Location | | | |
|---|---|---|---|---|---|---|
| | | | S | EV | PN | PE |
| Low level $N_2O$ | pH 4.5 | NCBI SRA No. | SRR22334937 | SRR22334946 | SRR22334945 | SRR22334938 |
| | | Number of reads | 52 255 324 | 63 252 210 | 61 708 064 | 69 517 460 |
| | pH 7.3 | NCBI SRA No. | SRR22334943 | SRR22334932 | SRR22334933 | SRR22334944 |
| | | Number of reads | 56 398 178 | 61 230 092 | 62 455 532 | 57 257 032 |
| High level $N_2O$ | pH 4.5 | NCBI SRA No. | SRR22334933 | SRR22334936 | SRR22334935 | SRR22334934 |
| | | Number of reads | 58 035 950 | 54 289 774 | 72 962 738 | 57 417 756 |
| | pH 7.3 | NCBI SRA No. | SRR22334939 | SRR22334942 | SRR22334941 | SRR22334940 |
| | | Number of reads | 60 836 792 | 47 671 962 | 57 093 952 | 58 052 954 |

**Table S3.** Metadata for published metagenomes downloaded from the European Nucleotide Archive.

| Biome | Location | pH | NCBI SRA No. | Number of samples | Reference |
|---|---|---|---|---|---|
| Temperate forest | British Columbia, Canada | 5.0-5.5 | ERR753925<br>ERR753926<br>ERR753927 | 3 | [4] |
| Tropical forest | Gigante Peninsula, Panama | 4.5 | SRR5262244<br>SRR5262250 | 2 | [5] |
| Temperate forest | New Hampshire, United States | 4.5 | SRR5580658<br>SRR5580692 | 2 | [6] |
| Tropical forest | Mount Kilimanjaro, Tanzania | 4.9-5.2 | ERR4660996<br>ERR4660997<br>ERR4660998<br>ERR4660999 | 4 | [7] |
| Agricultural land | Yunnan, China | 3.5-5.2 | SRR10260009<br>SRR10260010 | 2 | [8] |
| Temperate forest | Indiana, USA | 5.7 | SRR8437983<br>SRR7687007<br>SRR8437984<br>SRR7687014 | 4 | [9] |
| Permafrost | Canada | 5.5 | SRR1586253<br>SRR1586257<br>SRR1586264 | 3 | [10] |
| Agricultural land | Norway | 3.8-4.0 | ERR5023167<br>ERR5023168<br>ERR5023169 | 3 | [11] |
| [a]Tropical forest | Puerto Rico | 4.5 | Not available | 4 | [1] |
| Agricultural land | Illinois, USA | 6.1-7.5 | ERR1939172<br>ERR1939173<br>ERR1939174<br>ERR1939267<br>ERR1939269 | 5 | [12] |
| Tropical forest | Mount Kilimanjaro (RAU), Tanzania | 7.5 | ERR4661012<br>ERR4661013<br>ERR4661014 | 3 | [7] |

[a] The metagenomic datasets were deposited in the European Nucleotide Archive (ENA) under project PRJEB26500.

**Table S4.** Performance of microcosms established with tropical soils collected along an elevational gradient in the LEF and experiencing low versus high $N_2O$ concentrations and acidic versus circumneutral pH conditions.

| | | | EV | PN | PE | S |
|---|---|---|---|---|---|---|
| | | $N_2O$ | | | | |
| Low level $N_2O$ | pH 4.5 | # of feedings | 26 | 18 | 21 | 21 |
| | | Total amount (µmol) | 108 | 75 | 87 | 87 |
| | pH 7.3 | # of feedings | 34 | 26 | 29 | 31 |
| | | Total amount (µmol) | 141 | 108 | 120 | 129 |
| High level $N_2O$ | pH 4.5 | # of feedings | 8 | 8 | 5 | 7 |
| | | Total amount (µmol) | 3 328 | 3 328 | 2 080 | 2 912 |
| | pH 7.3 | # of feedings | 17 | 15 | 9 | 13 |
| | | Total amount (µmol) | 7 072 | 6 240 | 3 744 | 5 408 |

**Table S5.** Number of 16S rRNA gene sequences derived from the original soils and 16 $N_2O$-reducing microcosms.

| | **EV** | **PN** | **PE** | **S** |
|---|---|---|---|---|
| Original soil | 4 390 | 7 483 | 8 636 | 3 341 |
| pH4.5_0.1 | 18 442 | 11 514 | 13 988 | 59 035 |
| pH4.5_10 | 23 100 | 14 905 | 19 532 | 55 570 |
| pH7.3_0.1 | 12 235 | 16 713 | 12 631 | 23 729 |
| pH7.3_10 | 9 792 | 20 144 | 23 505 | 39 554 |

The numbers 0.1 and 10 reflect low (0.02 mM) and high (2 mM) $N_2O$ levels in the microcosms.

**Table S6.** Results of PERMANOVA based on weighted-UniFrac distance for the effects of pH and $N_2O$ on microbial community composition.

| **Factor** | **Df** | **Sum of Squares** | **$R^2$** | **F** | ***p*-value** |
|---|---|---|---|---|---|
| pH | 2 | 1.083 | 0.375 | 4.965 | 0.001 |
| $N_2O$ | 1 | 0.108 | 0.037 | 0.986 | 0.406 |
| pH × $N_2O$ | 1 | 0.059 | 0.020 | 0.539 | 0.861 |

**Table S7.** Number of *nosZ* reads identified in metagenome datasets derived from $N_2O$-reducing tropical soil microcosms using ROCker models.

| | Number of ***nosZ*** Sequences | | | | | | | |
|---|---|---|---|---|---|---|---|---|
| | **Clade I** | | | | **Clade II** | | | |
| | **Location** | | | | **Location** | | | |
| **Microcosm** | **S** | **EV** | **PN** | **PE** | **S** | **EV** | **PN** | **PE** |
| pH4.5_0.1 | 1 065 | 123 | 4 151 | 3 283 | 1 626 | 720 | 5 833 | 7 585 |
| pH4.5_10 | 8 | 105 | 16 506 | 245 | 69 | 3 479 | 22 940 | 3 529 |
| pH7.3_0.1 | 1 574 | 147 | 174 | 252 | 312 | 1 487 | 5 436 | 2 443 |
| pH7.3_10 | 6 140 | 58 | 2 913 | 171 | 31 604 | 474 | 10 420 | 7 165 |

The numbers 0.1 and 10 reflect low (0.02 mM) and high (2 mM) $N_2O$ levels in the microcosms.

**Table S8.** Summary statistics of the 17 high-quality MAGs derived from $N_2O$-reducing microcosms harboring *nosZ* genes.

| MAG ID [a] | Genome size (Mbp) | No. scaffolds | N50 (scaffolds) | GC (%) | No. CDS [b] | Completeness (%) | Contamination (%) |
|---|---|---|---|---|---|---|---|
| PN_pH4.5_0.1_MAG1 | 6.72 | 547 | 22792 | 68.6 | 6344 | 97.05 | 1.74 |
| PE_pH4.5_0.1_MAG2 | 6.39 | 623 | 15910 | 68.4 | 6109 | 90.22 | 3.01 |
| S_pH4.5_0.1_MAG3 | 6.31 | 1038 | 8619 | 68.1 | 6441 | 84.89 | 3.85 |
| EV_pH4.5_10_MAG4 | 5.42 | 139 | 97917 | 46.3 | 5146 | 99.43 | 2.69 |
| PN_pH4.5_10_MAG5 | 5.34 | 126 | 139753 | 46.4 | 5057 | 99.43 | 1.62 |
| PE_pH4.5_10_MAG6 | 4.62 | 107 | 98082 | 42.9 | 4372 | 100 | 1.28 |
| PE_pH4.5_10_MAG7 | 7.67 | 253 | 73205 | 51.6 | 6926 | 98.39 | 2.26 |
| PE_pH4.5_10_MAG8 | 2.55 | 570 | 6301 | 68.2 | 2774 | 84.03 | 2.36 |
| EV_pH7.3_0.1_MAG9 | 3.62 | 40 | 224397 | 63.8 | 3314 | 99.76 | 0.32 |
| PN_pH7.3_0.1_MAG10 | 3.69 | 42 | 405271 | 63.3 | 3363 | 99.76 | 0.95 |
| PE_pH7.3_0.1_MAG11 | 3.90 | 108 | 339534 | 63.1 | 3605 | 99.76 | 1.76 |
| S_pH7.3_0.1_MAG12 | 3.19 | 70 | 151307 | 62.5 | 3080 | 98.66 | 0.71 |
| EV_pH7.3_10_MAG13 | 5.34 | 183 | 70882 | 46.2 | 5090 | 84.18 | 2.17 |
| PN_pH7.3_10_MAG14 | 3.55 | 71 | 122344 | 63.5 | 3404 | 100 | 0.05 |
| PE_pH7.3_10_MAG15 | 3.45 | 44 | 115048 | 63.9 | 3281 | 98.34 | 0.43 |
| S_pH7.3_10_MAG16 | 3.58 | 82 | 72588 | 63.8 | 3321 | 98.48 | 0.32 |
| S_pH7.3_10_MAG17 | 6.80 | 86 | 206838 | 47.5 | 5053 | 98.89 | 3.13 |

[a] The numbers 0.1 and 10 reflect low (0.02 mM) and high (2 mM) $N_2O$ levels in the microcosms.

[b] CDS, coding sequence, is a region of DNA whose sequence codes for a protein.

**Table S9.** GTDB-Tk taxonomic classification of the taxa represented by the 17 high-quality MAGs harboring *nosZ* genes.

| MAG ID [a] | GTDB-Tk taxonomic classification |
|---|---|
| PN_pH4.5_0.1_MAG1 | d__Bacteria;p__Proteobacteria;c__Alphaproteobacteria;o__Rhizobiales;f__Xanthobacteraceae;g__Rhodoplanes;s__ |
| PE_pH4.5_0.1_MAG2 | d__Bacteria;p__Proteobacteria;c__Alphaproteobacteria;o__Rhizobiales;f__Xanthobacteraceae;g__Rhodoplanes;s__ |
| S_pH4.5_0.1_MAG3 | d__Bacteria;p__Proteobacteria;c__Alphaproteobacteria;o__Rhizobiales;f__Xanthobacteraceae;g__Rhodoplanes;s__ |
| EV_pH4.5_10_MAG4 | d__Bacteria;p__Firmicutes_B;c__Desulfitobacteriia;o__Desulfitobacteriales;f__Desulfitobacteriaceae;g__Desulfosporosinus;s__ |
| PN_pH4.5_10_MAG5 | d__Bacteria;p__Firmicutes_B;c__Desulfitobacteriia;o__Desulfitobacteriales;f__Desulfitobacteriaceae;g__Desulfosporosinus;s__ |
| PE_pH4.5_10_MAG6 | d__Bacteria;p__Firmicutes_B;c__Desulfitobacteriia;o__Desulfitobacteriales;f__Desulfitobacteriaceae;g__Desulfosporosinus;s__ |
| PE_pH4.5_10_MAG7 | d__Bacteria;p__Desulfobacterota;c__Desulfomonilia;o__Desulfomonilales;f__Desulfomonilaceae;g__Desulfomonile;s__ |
| PE_pH4.5_10_MAG8 | d__Bacteria;p__Actinobacteriota;c__Coriobacteriia;o__OPB41;f__PALSA-660;g__;s__ |
| EV_pH7.3_0.1_MAG9 | d__Bacteria;p__Proteobacteria;c__Gammaproteobacteria;o__Burkholderiales;f__Rhodocyclaceae;g__Azospira;s__ |
| PN_pH7.3_0.1_MAG10 | d__Bacteria;p__Proteobacteria;c__Gammaproteobacteria;o__Burkholderiales;f__Rhodocyclaceae;g__Azospira;s__ |
| PE_pH7.3_0.1_MAG11 | d__Bacteria;p__Proteobacteria;c__Gammaproteobacteria;o__Burkholderiales;f__Rhodocyclaceae;g__Azospira;s__ |
| S_pH7.3_0.1_MAG12 | d__Bacteria;p__Proteobacteria;c__Gammaproteobacteria;o__Burkholderiales;f__SulfuricEVlaceae;g__UBA2239;s__ |
| EV_pH7.3_10_MAG13 | d__Bacteria;p__Firmicutes_B;c__Desulfitobacteriia;o__Desulfitobacteriales;f__Desulfitobacteriaceae;g__Desulfosporosinus;s__ |
| PN_pH7.3_10_MAG14 | d__Bacteria;p__Proteobacteria;c__Gammaproteobacteria;o__Burkholderiales;f__Rhodocyclaceae;g__;s__ |
| PE_pH7.3_10_MAG15 | d__Bacteria;p__Proteobacteria;c__Gammaproteobacteria;o__Burkholderiales;f__Rhodocyclaceae;g__Azospira;s__ |
| S_pH7.3_10_MAG16 | d__Bacteria;p__Proteobacteria;c__Gammaproteobacteria;o__Burkholderiales;f__Rhodocyclaceae;g__Azospira;s__ |
| S_pH7.3_10_MAG17 | d__Bacteria;p__Firmicutes_B;c__Desulfitobacteriia;o__Desulfitobacteriales;f__Desulfitobacteriaceae;g__Desulfitobacterium;s__ |

[a] The numbers 0.1 and 10 reflect low (0.02 mM) and high (2 mM) $N_2O$ levels in the microcosms.

**Table S10.** Analysis of 17 high-quality MAGs harboring *nosZ* genes using the Microbial Genome Atlas (MiGA). Listed for each MAG is the average amino acid identity (AAI), the closest relative, and the novelty of each MAG. The taxonomic novelty is determined by the maximum AAI value compared to genomes in the TypeMat database. The p-value is estimated from the empirical distribution observed in all reference genomes in the NCBI Reference Sequence Database (RefSeq) at each taxonomic level and indicates the probability of the observed AAI between genomes of the same taxon. The TypeMat database contains assemblies from type materials in Archaea and Bacteria (as flagged by NCBI) including both complete and draft genomes.

| MAG ID [a] | Best classification level | AAI | Closest relative | Novelty |
|---|---|---|---|---|
| PN_pH4.5_0.1_MAG1 | **Order** *Rhizobiales* (p-value: 0.0024) | 71.28 | *Rhodoplanes roseus* | **Species** (p-value: 0.00252) |
| PE_pH4.5_0.1_MAG2 | **Order** *Rhizobiales* (p-value: 0.0024) | 71.17 | *Rhodoplanes roseus* | **Species** (p-value: 0.00252) |
| S_pH4.5_0.1_MAG3 | **Order** *Rhizobiales* (p-value: 0.0024) | 70.32 | *Rhodoplanes roseus* | **Species** (p-value: 0.00252) |
| EV_pH4.5_10_MAG4 | **Family** *Peptococcaceae* (p-value: 0.0057) | 78.82 | *Desulfosporosinus orientis* DSM 765 | **Species** (p-value: 0.00269) |
| PN_pH4.5_10_MAG5 | **Family** *Peptococcaceae* (p-value: 0.0057) | 79.2 | *Desulfosporosinus orientis* DSM 765 | **Species** (p-value: 0.00269) |
| PE_pH4.5_10_MAG6 | **Family** *Peptococcaceae* (p-value: 0.0061) | 77.84 | *Desulfosporosinus lacus* DSM 15449 | **Species** (p-value: 0.00257) |
| PE_pH4.5_10_MAG7 | **Phylum** Proteobacteria (p-value: 0.0011) | 45.04 | *Desulfosoma caldarium* | **Family** (p-value: 0.0151) |
| PE_pH4.5_10_MAG8 | **Phylum** Actinobacteria (p-value: 0.0011) | 44.04 | *Rhabdothermincola sediminis* | **Family** (p-value: 0.0119) |
| EV_pH7.3_0.1_MAG9 | **Order** *Rhodocyclales* (p-value: 0.0024) | 69.99 | *Azospira oryzae* | **Species** (p-value: 0.00252) |
| PN_pH7.3_0.1_MAG10 | **Order** *Rhodocyclales* (p-value: 0.0024) | 69.98 | *Azospira oryzae* | **Species** (p-value: 0.00252) |
| PE_pH7.3_0.1_MAG11 | **Order** *Rhodocyclales* (p-value: 0.0024) | 69.99 | *Azospira oryzae* | **Species** (p-value: 0.00252) |
| S_pH7.3_0.1_MAG12 | **Class** *Betaproteobacteria* (p-value: 0.00073) | 56.82 | *Sulfurimicrobium lacus* NZ AP022853 | **Species** (p-value: 0.000457) |
| EV_pH7.3_10_MAG13 | **Family** *Peptococcaceae* (p-value: 0.0057) | 79.42 | *Sulfurimicrobium lacus* NZ AP022853 | **Species** (p-value: 0.00269) |
| PN_pH7.3_10_MAG14 | **Order** *Rhodocyclales* (p-value: 0.0021) | 66.86 | *Dechloromonas hortensis* | **Species** (p-value: 0.002) |
| PE_pH7.3_10_MAG15 | **Order** *Rhodocyclales* (p-value: 0.0024) | 69.21 | *Azospira oryzae* | **Species** (p-value: 0.00252) |
| S_pH7.3_10_MAG16 | **Order** *Rhodocyclales* (p-value: 0.0024) | 69.64 | *Azospira oryzae* | **Species** (p-value: 0.00252) |
| S_pH7.3_10_MAG17 | **Genus** *Desulfitobacterium* (p-value: 0.0079) | 89.81 | *Desulfitobacterium chlororespirans* DSM 11544 | **Subspecies** (p-value: 0.000182) |

[a] The numbers 0.1 and 10 reflect low (0.02 mM) and high (2 mM) $N_2O$ levels in the microcosms.

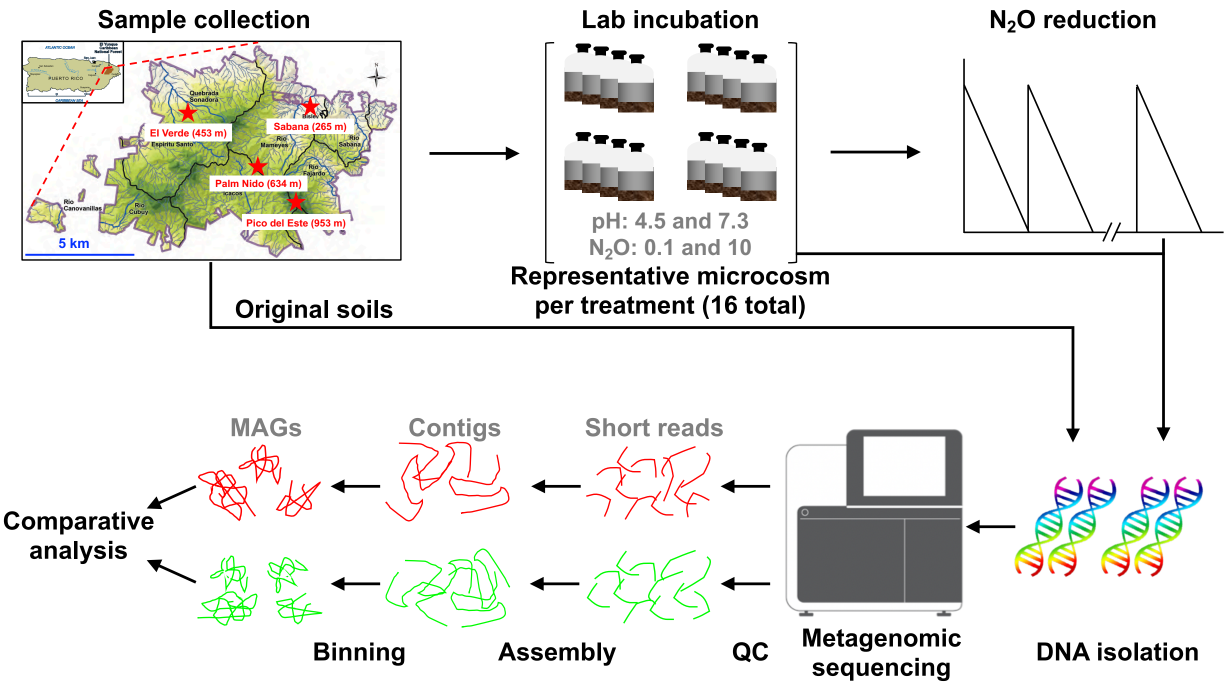


**Figure S1.** Schematic of the study site and overview of the experimental design. The numbers 0.1 and 10 represent the low and high levels of $N_2O$ maintained in the microcosms, respectively. Microcosms had two technical replicates, which showed similar performance, and one microcosm per treatment was selected for metagenome sequencing (16 total). Negative controls included heat-killed (autoclaved) replicates and microcosms without $N_2O$ but with lactate at pH 4.5 for each soil sample. The metagenomes of the four original soils have been analyzed in a prior study [1] and are available in the European Nucleotide Archive under project PRJEB26500.

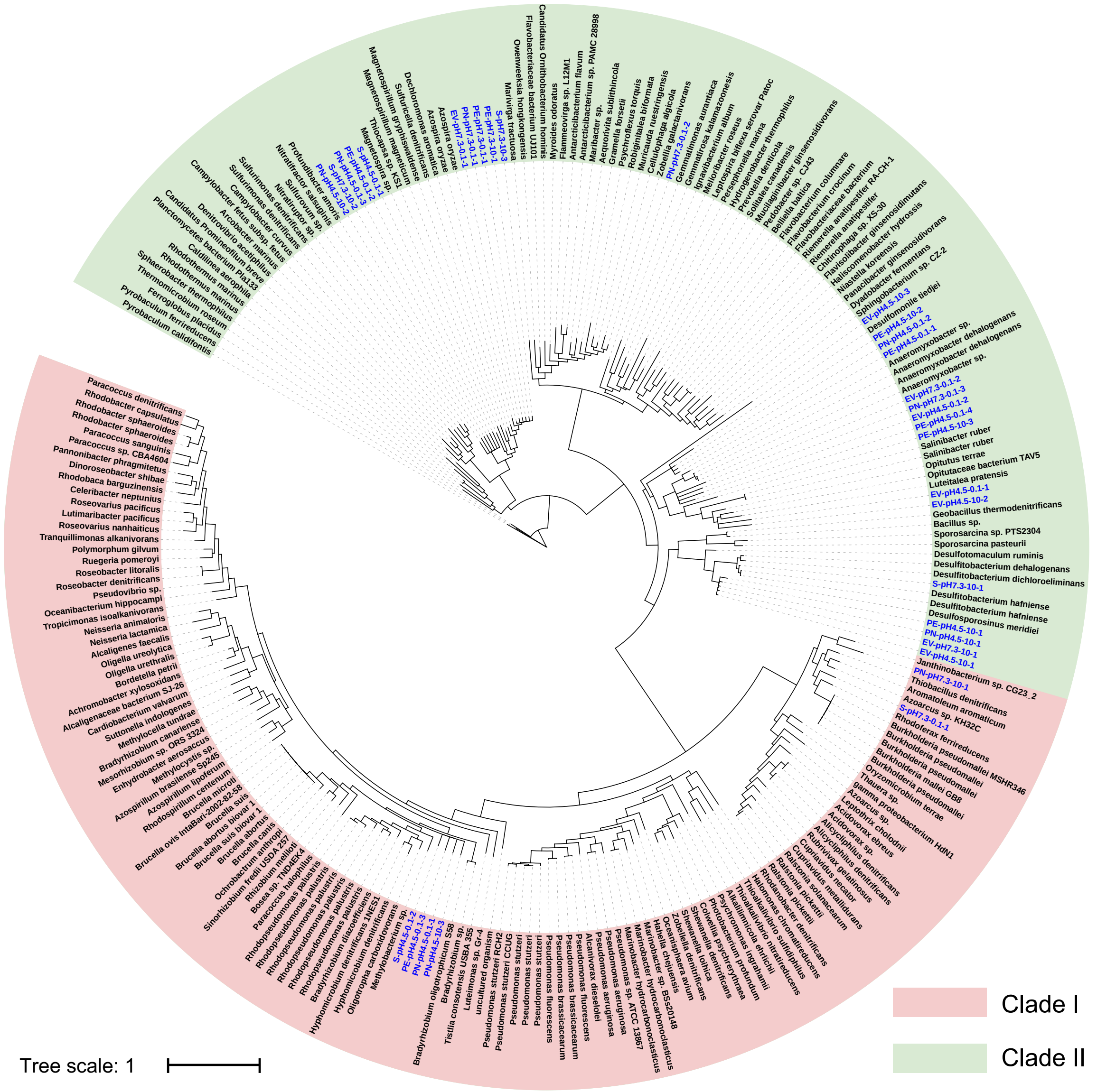


**Figure S2.** Phylogenetic diversity of the putative near full-length *nosZ* sequences recovered from the assembled contigs and *nosZ* from the reference *nosZ* database. The red half circle indicates clade I *nosZ* and the green half circle depicts clade II *nosZ*. Branch labels in blue represent the *nosZ* contigs assembled from the metagenomic data. The numbers 0.1 and 10 reflect low (0.02 mM) and high (2 mM) $N_2O$ levels in the microcosms.

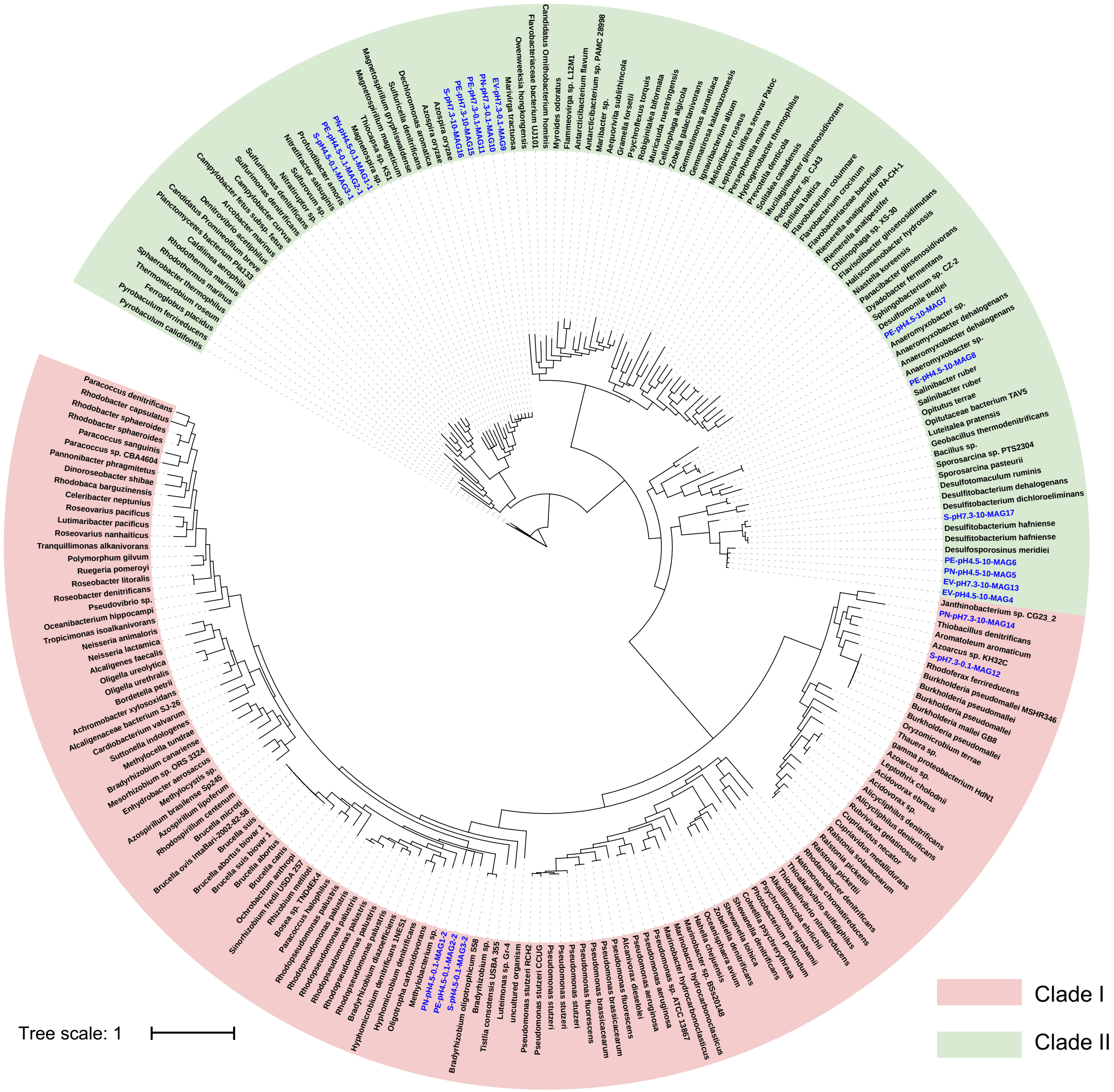


**Figure S3.** Phylogenetic diversity of the near full-length *nosZ* sequences recovered from the high-quality MAGs and *nosZ* from the reference *nosZ* database. The red and the green half circles indicate clade I *nosZ* and clade II *nosZ*, respectively. Branch labels in blue indicate *nosZ* sequences from the MAGs. The numbers 0.1 and 10 reflect low (0.02 mM) and high (2 mM) $N_2O$ levels in the microcosms.

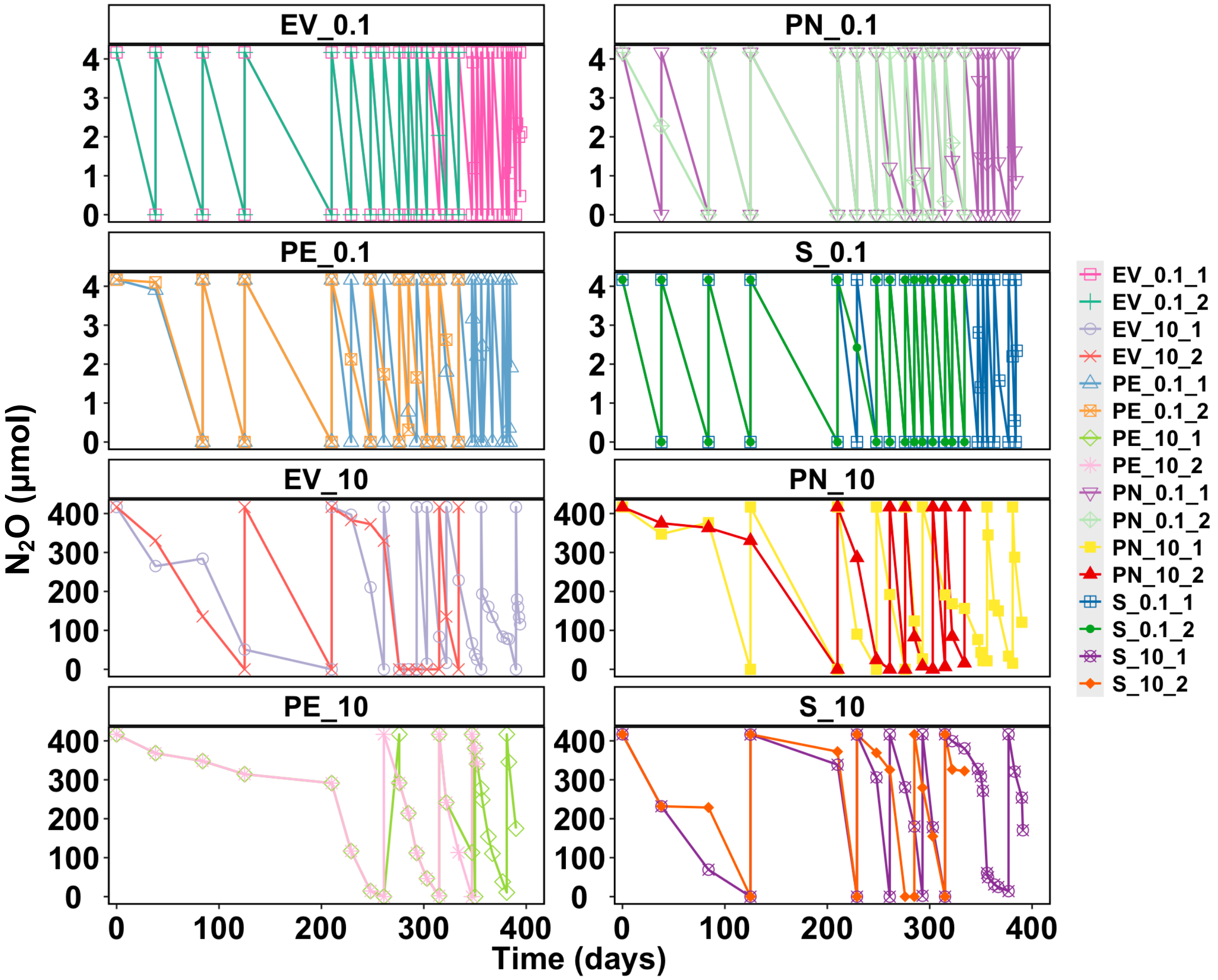


**Figure S4.** $N_2O$ reduction profiles in acidic (pH 4.5) tropical soil microcosms. The numbers 0.1 and 10 reflect low (0.02 mM) and high (2 mM) $N_2O$ levels in the microcosms. Replicate microcosms showed similar performance.

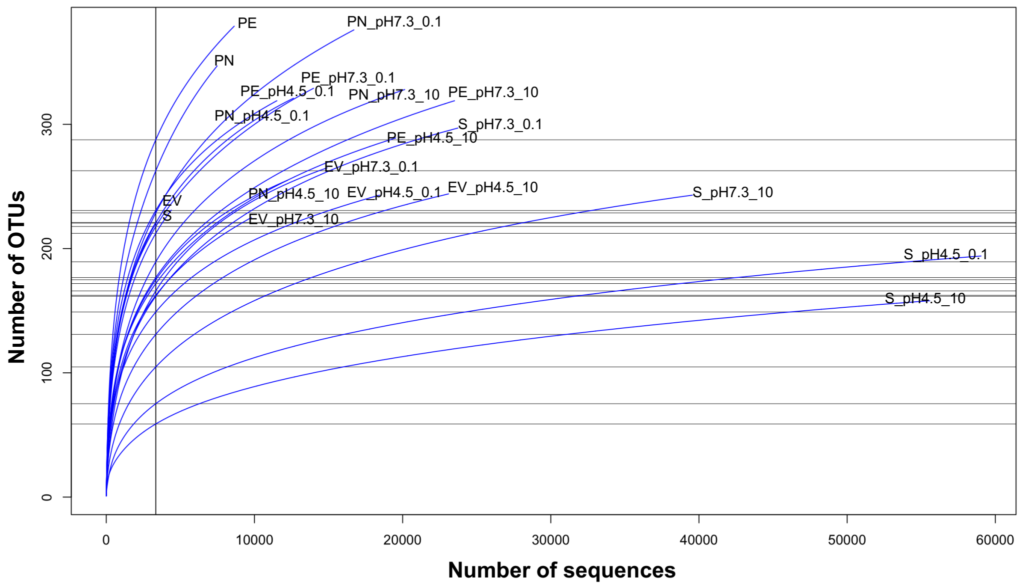


**Figure S5.** Rarefaction curves of 16S rRNA gene sequences recovered from metagenomic datasets. Curves represent the number of unique OTUs recovered, defined at the 97% nucleotide sequence identity level, for the number of sequences analyzed, and reflect the extent of OTU diversity within the samples and what fraction of this diversity was sampled.

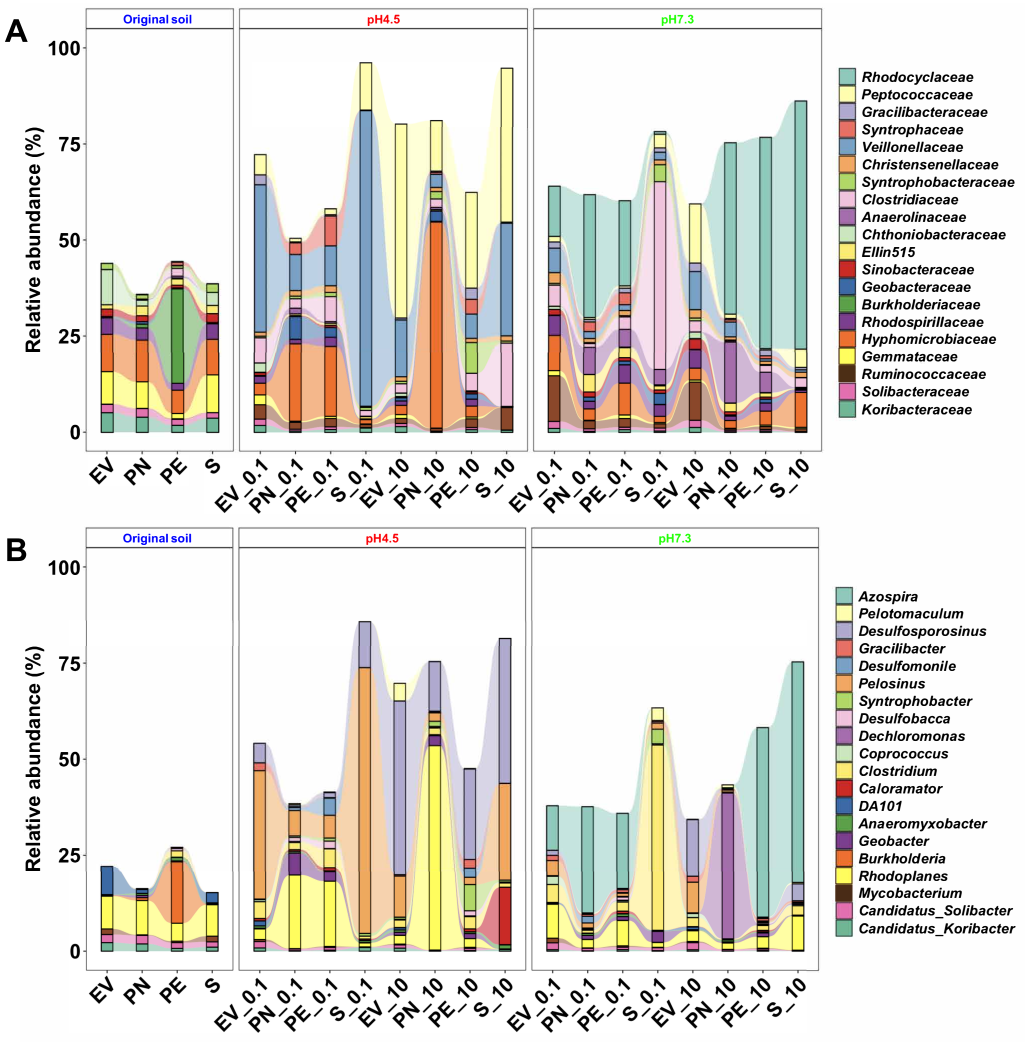


**Figure S6.** Microbial community compositions based on 16S rRNA gene fragments recovered from metagenome data from original LEF soils and the $N_2O$-reducing microcosms maintained at pH 4.5 and pH 7.3 and with low (0.02 mM) and high (2 mM) levels of $N_2O$. (A) Relative abundances of the top 20 families in the original soils and the microcosms maintained under the different enrichment conditions. (B) Relative abundances of the top 20 genera in the original soils and the microcosms maintained under the different enrichment conditions. The numbers 0.1 and 10 reflect low (0.02 mM) and high (2 mM) $N_2O$ levels in the microcosms.

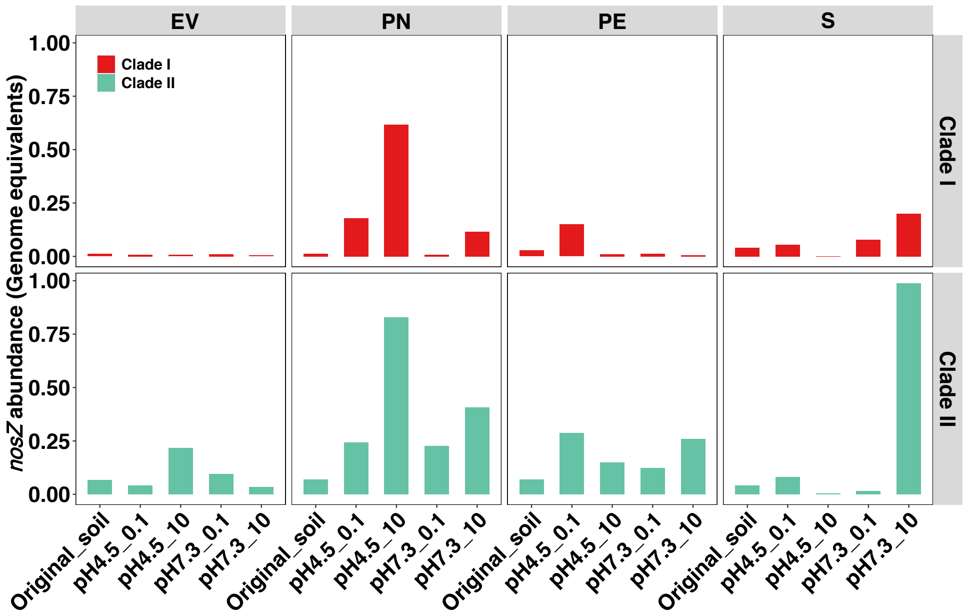


**Figure S7.** Abundance of *nosZ* genes in the original LEF soils and the 16 $N_2O$-reducing microcosms selected for metagenome sequencing. The numbers 0.1 and 10 reflect low (0.02 mM) and high (2 mM) $N_2O$ levels in the microcosms.

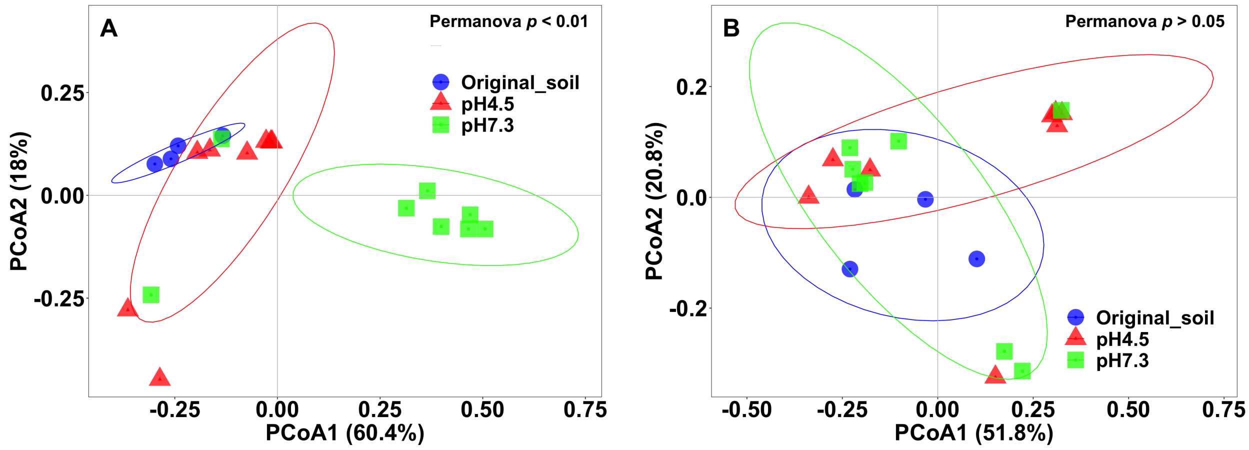


**Figure S8.** Differences in microbial community composition between the original soils and the respective $N_2O$-reducing microcosms based on weighted Unifrac analysis of *nosZ* gene fragments recovered in the metagenomes using ROCker. (A) Beta diversity based on clade II *nosZ* sequences. (B) Beta diversity based on clade I *nosZ* sequences $N_2O$. Samples are visualized by principal coordinate analysis (PCoA) with colors representing the original soil (blue) and pH (pH 4.5, red; pH 7.3, green). The ellipses represent the 95% confidence intervals.

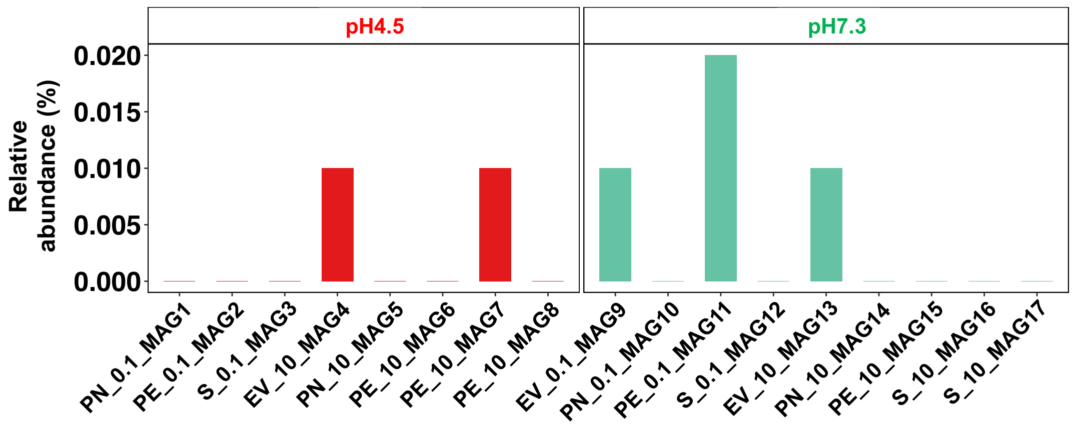


**Figure S9.** The relative abundance of each MAG harboring a *nosZ* gene derived from $N_2O$-reducing microcosms in the corresponding original soils based on metagenomic reads competitively mapped against the MAGs.

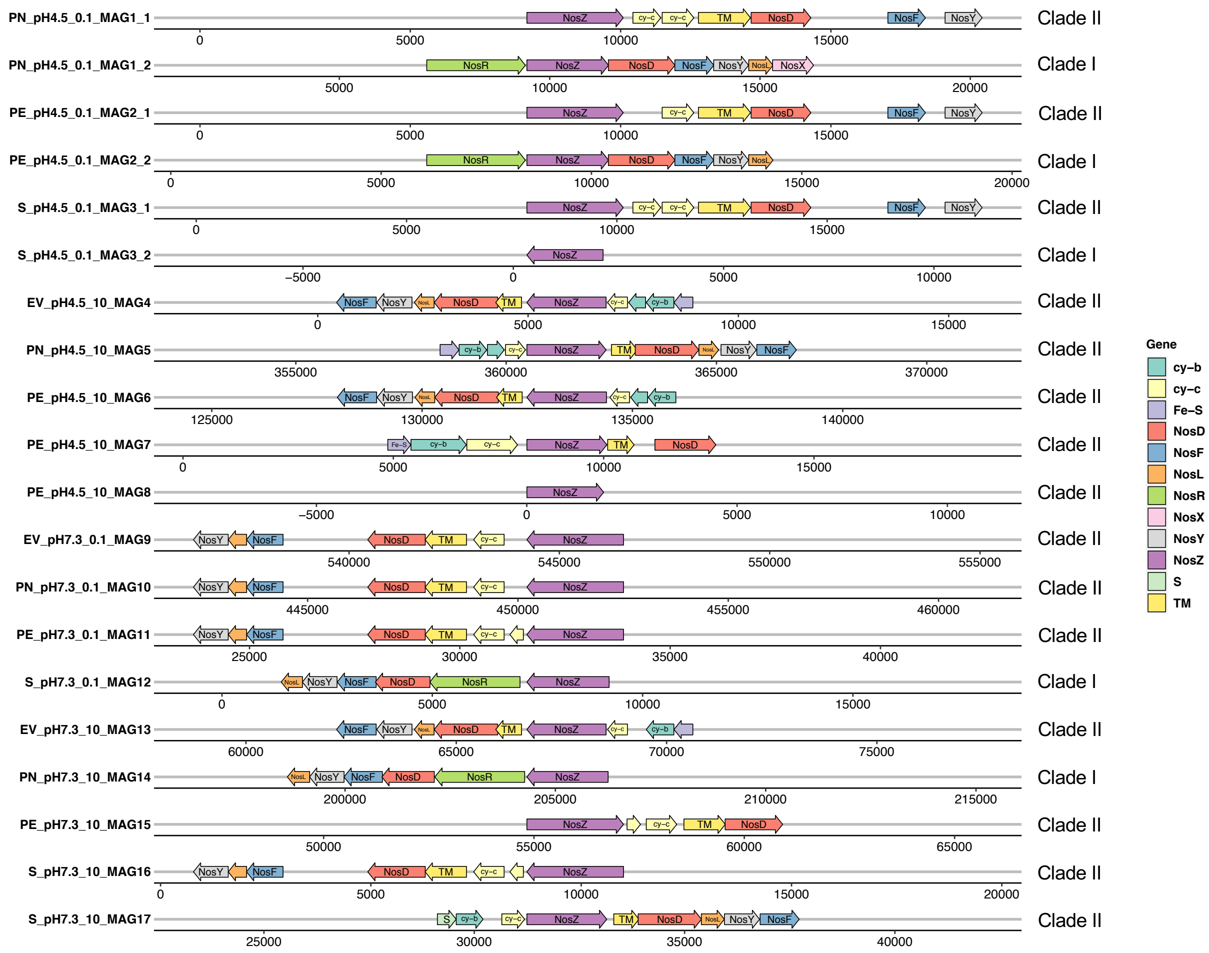


**Figure S10.** Comparison of *nos* clusters in the 17 MAGs harboring *nosZ* genes derived from $N_2O$-reducing microcosms. *nos* cluster genes were annotated using the SEED subsystem and validated by using the BLAST. Transmembrane helices in transmembrane proteins were identified using the TMHMM2 (http://smart.embl-heidelberg.de/). cy-b and cy-c represent b-type and c-type cytochromes, respectively. Fe-S, S, and TM represent genes encoding iron-sulfur-binding proteins, Rieske iron-sulfur proteins, and transmembrane proteins, respectively. The numbers 0.1 and 10 reflect low (0.02 mM) and high (2 mM) $N_2O$ levels in the microcosms. The numbers 1 and 2 after the last low dash in the left label represent the first and second *nosZ* genes identified in the corresponding MAG.

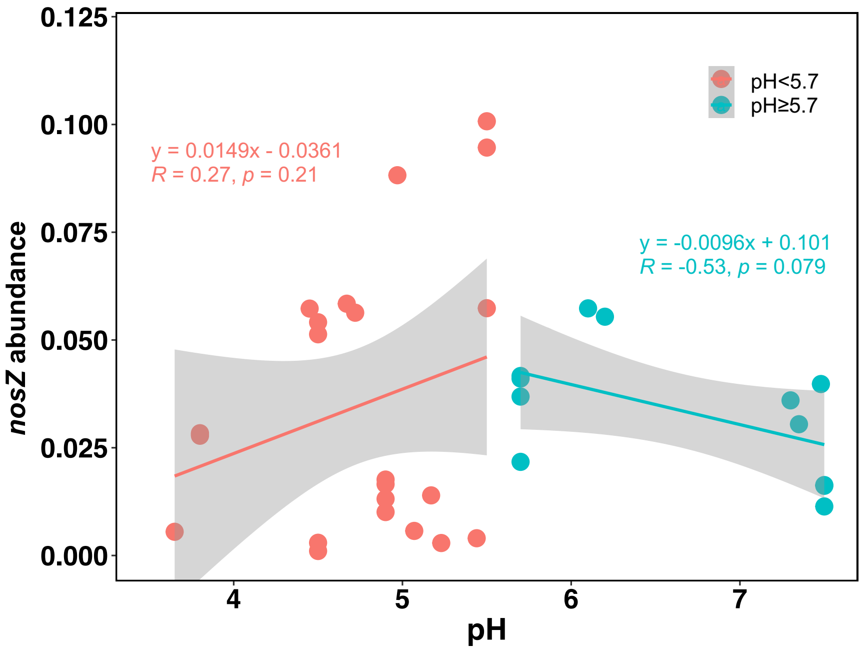


**Figure S11.** Relationships between pH and the total abundance of *nosZ* genes per genome equivalent. Detailed information about the metagenome datasets is provided in Table S3. *nosZ* genes were searched against a customized database of *nosZ* based on near full-length *nosZ* identified in the acidic $N_2O$-reducing microcosms. The plots show the total abundance of *nosZ* genes per genome equivalent. Gray areas represent 95% confidence intervals.
